# CRISPR-Detector: Fast and Accurate Detection, Visualization, and Annotation of Genome-Wide Mutations Induced by Gene Editing Events

**DOI:** 10.1101/2022.02.16.480781

**Authors:** Lei Huang, Dan Wang, Haodong Chen, Jinnan Hu, Xuechen Dai, Chuan Liu, Anduo Li, Xuechun Shen, Chen Qi, Haixi Sun, Dengwei Zhang, Tong Chen, Yuan Jiang

## Abstract

The leading edge of genome editing, driven by CRISPR/Cas technology, is revolutionizing biotechnologies. In pace with the burst of novel genome editing methods, bioinformatic tools to monitor the on/off-target events need to be more evolved as well. Existing tools suffer from limitations in speed and scalability, and they especially struggle with whole genome sequencing (WGS) data analysis which has the potential to detect off-target mutations at a genome-wide level via an unbiased manner. Here, we introduce our CRISPR-detector tool, which is a web-hosted or locally deployable pipeline with five key innovations: 1) optimized scalability allowing for WGS data analysis beyond BED (Browser Extensible Data) file-defined regions; 2) improved accuracy benefiting from haplotype-based variant calling to handle sequencing errors; 3) treated and control sample co-analysis to remove background variants existing prior to genome editing; 4) integrated structural variation (SV) calling; and 5) functional and clinical annotation of editing-induced mutations.

## Introduction

The clustered regularly interspaced short palindromic repeats (CRISPR)-based technologies, which are inspired by adaptive immune systems in microorganisms, have revolutionized the field of genome engineering. The RNA-guided CRISPR-associated (Cas) nuclease is used for genome editing by simply specifying a ∼20-nt targeting sequence within its guide RNA (Jinek et al., 2012). Since the first report of CRISPR/Cas9 editing technology in 2012, various novel CRISPR-based genome editing tools have been introduced by different groups (Cong et al., 2013; Zetsche et al., 2015; Abudayyeh et al., 2016; Liu et al., 2019; Pausch et al., 2020). Genome editing methods have progressively become more flexible and precise, including innovations such as base editors, which enable the direct conversion of one nucleotide to another without producing DSBs in the target sequence, and prime editing which can create targeted indels and introduce all 12 types of point mutations (Komor et al., 2016; Gaudelli et al 2017; Kim et al., 2017; Anzalone et al 2019). This technology creates exciting new opportunities for a plethora of life science applications including therapeutic treatment, drug discovery, biofuel production, agriculture, and more. Early clinical trials demonstrated CRISPR’s general safety and utility, and many more trials are underway (Lu et al., 2020; Frangoul et al., 2021; Gillmore et al., 2021). Despite the many important applications of these editing tools, they each have their own limitations. One common concern of the CRISPR/Cas technology is that editing events may occur at undesired locations, i.e., off-target events. As a result, the increased adoption of CRISPR-based treatment makes precision analysis—accurately detecting on- and off-target events—a top priority for regulators including the FDA. Accordingly, the FDA published guidance related to genome editing products in March 2022 (https://www.fda.gov/regulatory-information/search-fda-guidance-documents/human-gene-therapy-products-incorporating-human-genome-editing), in which they raised concerns multiple times about detecting the type and frequency of gene editing events. Therefore, the bioinformatic tools to monitor on/off-target events need to become more accurate and capable of detecting true editing-induced mutations—mutations introduced during gene editing as opposed to germline or previously existing somatic variants.

Targeted amplicon sequencing has become the gold standard for the validation of genome editing experiments (Akcakaya et al., 2018). Sequencing of amplicon focuses on specific genomic areas at high depth, making it suitable to detect low allele frequency (AF) mutations. Potential off-target sites can be predicted using tools such as Cas-OFFinder to search the on-target DNA sequences against the whole genome while allowing mismatches (preferred value: 1-5) and gaps (preferred value: 0-2) (Bae et al., 2014). Primers are then designed to flank each on/off-target cut-site and pooled for multiplex amplification assays. Genomic DNA from each edited sample is transformed into libraries and enriched via polymerase chain reaction (PCR) for target enrichment. Amplicons are then purified and sequenced using NGS platforms. NGS technologies offer ultra-deep sequencing of amplicons ranging from 100 to 500 bp.

Along with the development of targeted amplicon sequencing technologies, tools to monitor on/off-target events have been developed. Examples include CRISPR genome analyzer (CRISPR-GA) (Güell et al., 2014), AGEseq (Xue and Tsai, 2015), ampliconDIVider (Varshney et al., 2015), Cas-Analyzer (Park et al., 2017), ampliCan (Labun et al., 2019), and CRISPResso1/2 (Pinello et al., 2016; Clement et al., 2019). However, there are four main limitations to existing tools. The first involves speed and scalability; most tools are designed to consider single or only a few amplicons for analysis. However, with the increasing need to comprehensively cover all potential off-target sites, large panel and even WGS analysis serve as good options. Second, tools including CRISPR-GA, AGEseq, ampliconDIVider, Cas-Analyzer, and CRISPResso1/2 do not support the co-analysis of paired treatment/control sequencing data, and thus previously existing single nucleotide polymorphisms (SNPs) and indels could be interpreted as CRISPR/Cas-induced mutations and lead to the incorrect quantification of mutation events. Third, besides SNPs and indels, structural variation is generally defined as an alteration within a region of DNA approximately 1 kb to 3 Mb in size (the operational range of structural variants has widened to include events > 50 bp) including deletions, duplications, copy-number variants, insertions, inversions, and translocations, which is ignored by current analysis. The fourth drawback is that most existing pipelines do not estimate the functional or clinical consequences resulting from editing-induced mutations. This annotation is quite important for researchers whose genome editing projects are for therapeutic purposes. To overcome these limitations, we herein propose a one-stop CRISPR-detector platform.

## Results

### Design of CRISPR-detector

CRISPR-detector is designed to be able to process large datasets such as WGS data and output accuracy results. The pipeline accepts individual original FASTQ sequencing files from respective CRISPR/Cas-treatment and control experiments. Each input FASTQ file includes sequencing data from one or multiple amplicons, e.g., one on-target and its putative off-target amplicons. Then pipeline processes data in three procedures: Pre-processing step to align and sort reads; Variants Identification step to call and filter SNPs, indels, and SVs that could be resulted from editing events; and Output step to annotate and visualize results (Fig. 1A).

**Fig. 1.**
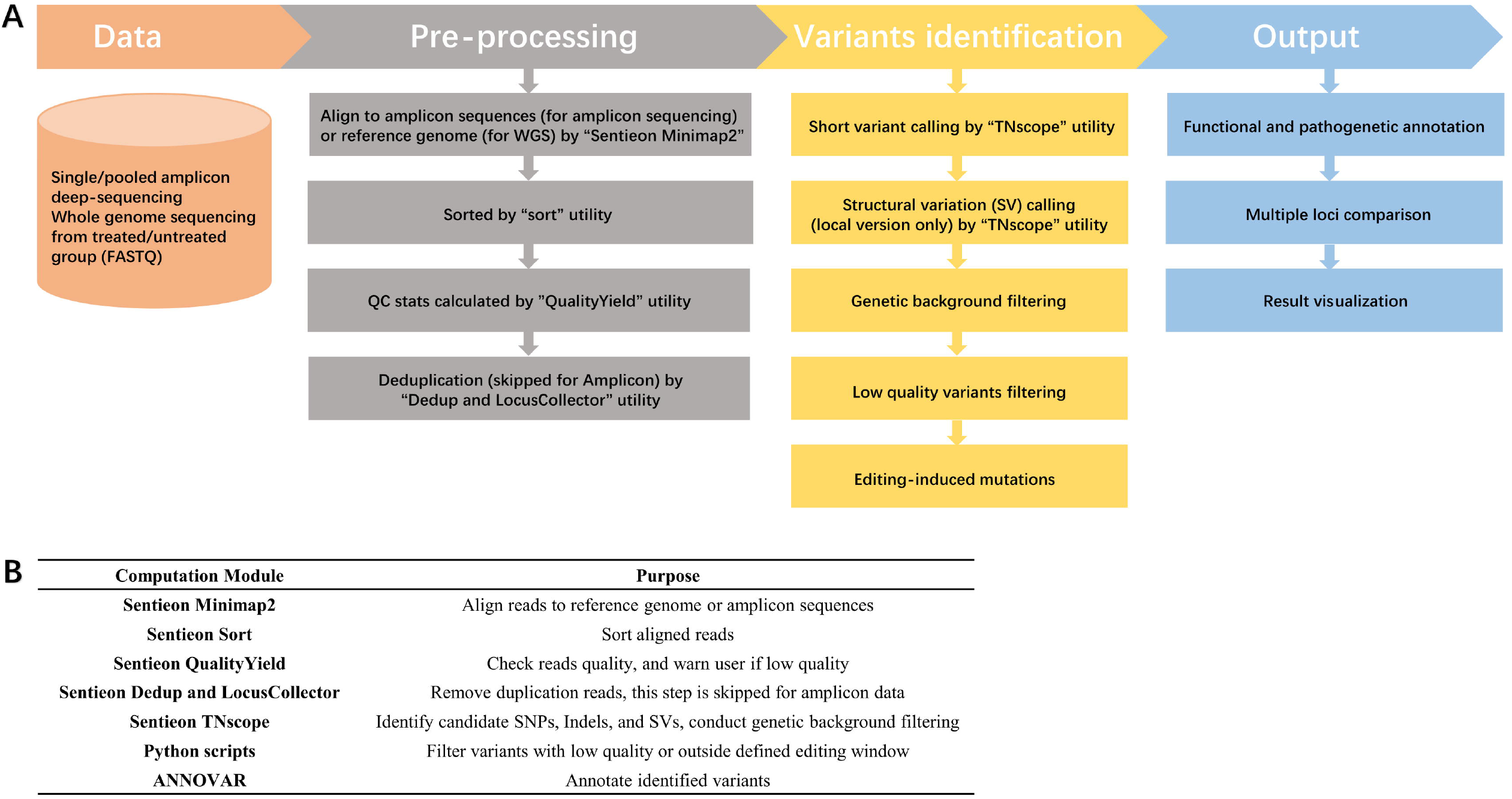
Flowchart and module description of CRISPR-detector. **A:** CRISPR-detector accepts single/pooled amplicon deep-sequencing as well as WGS data as input. The control group sample is utilized for removing background variants existing prior to genome editing for the treatment group sample. CRISPR-detector outputs visualization results and functional annotations of editing-induced mutations for targeted amplicon sequencing data, and VCF format text file for WGS data. **B:** Description of analysis modules that have been used in the pipeline.

During its Pre-processing and variant identification steps, CRISPR-detector chooses to utilize Sentieon high performance computation modules to reach higher speed which is necessary for WGS analysis (Fig. 1B). Sentieon’s modules are 3-20x times faster than corresponding opensource tools, and have better scalability for high threads computation environment. The Sentieon acceleration comes from a complete rewrite of the mathematical models of the open-source tools, using only C and C++ highly efficient languages, and better software engineering and resource management system to fully utilize CPU and memory resources.

The output of the web-hosted version of CRISPR-detector includes comprehensive analysis, intuitive visualization, and locus comparison. Specifically, CRISPR-detector primarily outputs: i) a summary webpage comparing and visualizing editing outcomes of the input amplicons; ii) detailed information on editing outcomes including frequency of SNPs and indels and a histogram of the indel sizes across the reference amplicon in the quantification window; iii) annotation of editing-induced mutations showing whether encoded protein sequences will be changed with filter-based annotation of mutations of clinical significance sourced from the ClinVar database (Landrum et al., 2018), including pathogenic mutations and those related to drug response, histocompatibility, and more.

In addition to the web-hosted version, a locally deployable CRISPR-detector pipeline focusing on WGS analysis and SV calling was developed. The pipeline calls not only regular SVs caused by the editing process but also viral vector insertion from virus-mediated gene editing. The whole genome analysis capability provided by the locally deployable version is unique and not available in any other gene editing analysis pipelines.

### Performance Benchmarks

#### 1. SNP/Indel Identification from Simulated Amplicon Datasets

The accurate detection of SNPs and indels is one of the key features of a gene editing analysis pipeline. To benchmark the accuracy of CRISPR-detector, we generated simulated testing reads representing actual sequencing data and compared the accuracy with three other popular pipelines including Cas-Analyzer, AGEseq, and CRISPResso2. We introduced SNPs (1-3 per read) and indels (5-50 bp length) into the simulated 150-bp paired-end reads at 50% frequency together with errors introduced from the MiSeq default profile provided by the ART simulation tool (Huang et al., 2012). The average running time was calculated from 3 repeated runs. The speed of Cas-Analyzer and AGEseq are not shown as they were tested on different computation platforms.

Compared to CRISPResso2, CRISPR-detector showed seven times faster processing speed, and the reported mutation rate was closer to the expected value of 0% for the control dataset and 50% for all other datasets (Fig. 2A). The faster speed of CRISPR-detector benefits from utilizing Sentieon high performance computation modules, such as alignment, sort, and variant calling. It should be noted that CRISPR-detector’s variant calling pipeline is based on haplotype assembly, which eliminates sequencing errors more easily compared with simple pileup-based read statistics.

**Fig. 2.**
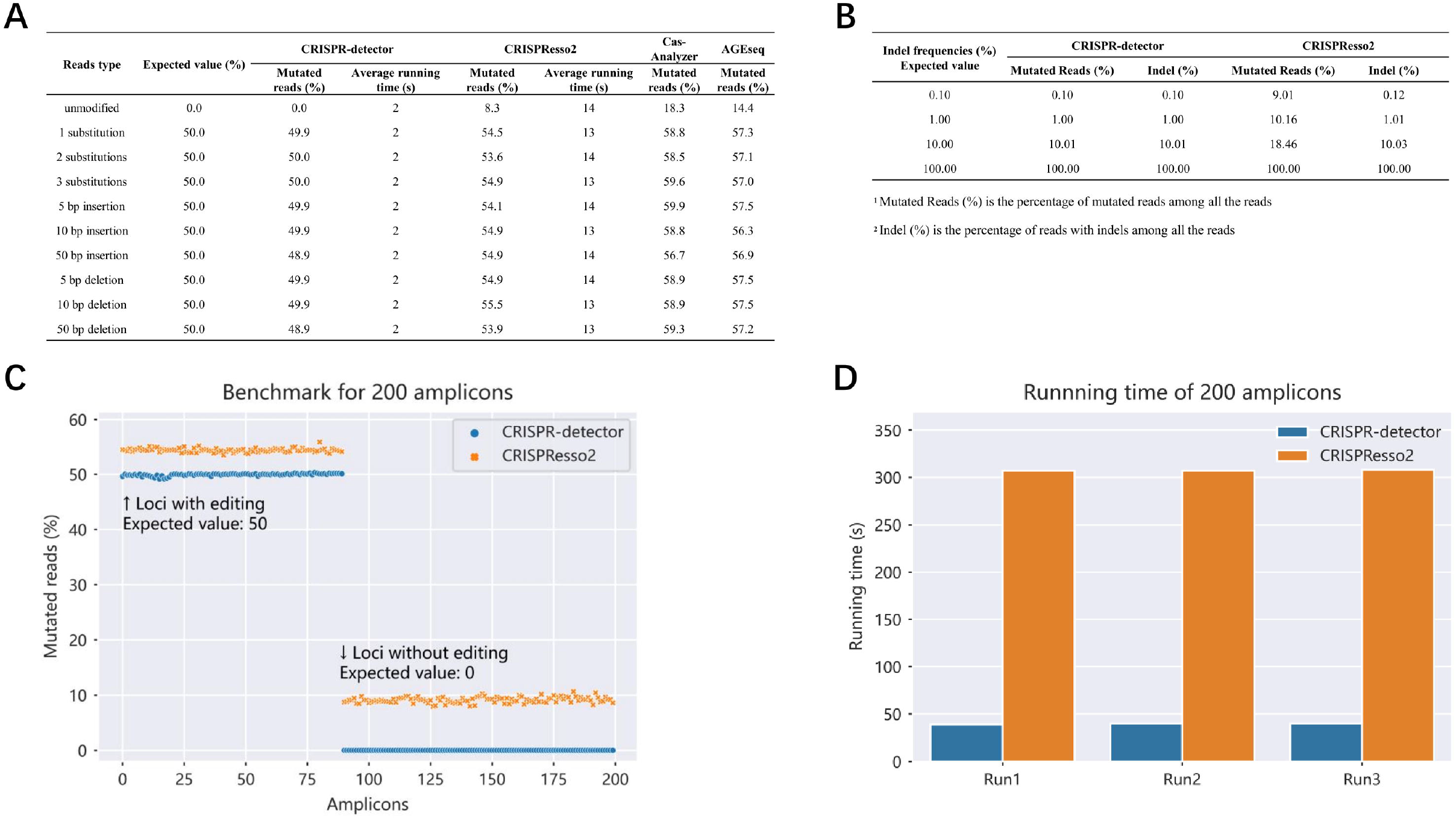
Accuracy and running time benchmark for single amplicon and 200 amplicons dataset. Amplicons in the format of 150 bp paired-end reads were simulated, to represent a large panel in which SNPs, indels, and sequencing errors (profiled by ART) were introduced. **A:** Simulated SNPs (1-3 per read) and indels (5-50 bp length) at 50% frequency were inserted. Compared to CRISPResso2, CRISPR-detector showed 7 times faster processing speed and reported mutated reads % closer to the expected value. **B:** 5 bp indel was introduced at multiple frequencies ladder ranging 0.01 % to 100 %, and processed by CRISPR-detector and CRISPResso2. Although neither tool correctly reported indel frequency at 0.01% level, CRISPR-detector outperformed CRISPResso2 at 0.1%-10% indels detection accuracy, by returning both mutated reads (%) and Indel (%) closer to expected values. **C:** CRISPR-detector outperforms CRISPResso2 on all 200 amplicons tested as its reported mutated reads frequency were closer to the expected 50% and 0% value, while CRISPResso2 overestimated mutated reads frequency by about 5%. **D:** CRISPR-detector runs 8 times faster than CRISPResso2 on these 200 amplicons dataset, slightly faster than the acceleration rate observed from the single amplicon benchmark.

We then performed more detailed variant detection testing to evaluate CRISPR-detector’s performance on indels at different frequencies and lengths. We designed a 5-bp indel in a single simulated amplicon dataset at 5 different frequencies starting from 0.01%. The testing dataset was processed by CRISPR-detector and CRISPResso2. CRISPR-detector reported nearly the exact expected frequency value, whereas CRISPResso2 included many more reads with sequencing errors incorrectly reported as mutated reads (Fig. 2B). TNscope (Freed et al., 2018), the variant calling module of CRISPR-detector performs well on 0.1% AF variants, because it utilizes a haplotype assembly method to filter out sequencing error, and this strategy makes it possible to filter out sequencing errors and identify very low AF variants as low as 0.1%.

Next, we simulated additional datasets to test the limits of CRISPR-detector in detecting indels of different lengths. Based on the general perception of the length limitations caused by short read sequencing, CRISPR-detector’s indel detection length is set as 50 bp for insertion and 71 bp for deletion. To demonstrate this performance, we simulated 50 amplicon datasets with insertions ranging from 1-50 bp, and 72 amplicons with deletions ranging from 1-72 bp, each at 50% frequency. The frequencies reported by CRISPR-detector across 1-50 bp insertions and 1-71 bp deletions aligned very close to the expected value. In comparison, CRISPResso2’s reported value was higher than expected, and it failed to detect deletions larger than 50 bp (Fig. S1). For insertions and deletions longer than 50 bp, CRISPR-detector will identify them through structural variation analysis modules.

CRISPR-detector can perform co-analysis with control sample data as input to address existing germline mutations and possible somatic or hematopoiesis-related mutations that were not introduced by a gene editing event (García-Nieto et al., 2019). In this simulated co-analysis testing dataset, some of the variants were inserted in both treatment and control reads to benchmark the pipeline’s ability to detect intrinsic “true positive” mutations (1-bp insertion at position 105 bp) in a noisy background with 7% sequencing error, the same setting used for CRISPResso2. For all read types except the 50-bp insertion, CRISPR-detector successfully identified true editing events without false positives from variants existing in control samples (Table 1).

**Table 1.** Identification of simulated insertion using co-analysis dataset.

| Simulated Reads Type |  |  | Variant Detected |  |  |  |
| --- | --- | --- | --- | --- | --- | --- |
| Control Reads Type | Inserted Reads Type | Expected Value (%) | POS | REF | ALT | Mutated reads (%) |
| 1 substitution | 1 substitution & 1 bp insertion | 50.0 | 105 | - | C | 49.6 |
| 2 substitutions | 2 substitutions & 1 bp insertion | 50.0 | 105 | - | C | 47.9 |
| 3 substitutions | 3 substitutions & 1 bp insertion | 50.0 | 105 | - | C | 47.9 |
| 5 bp deletion | 5 bp deletion & 1 bp insertion | 50.0 | 105 | - | C | 49.8 |
| 10 bp deletion | 10 bp deletion & 1 bp insertion | 50.0 | 105 | - | C | 65.8 |
| 50 bp deletion | 50 bp deletion & 1 bp insertion | 50.0 | 105 | - | C | 64.0 |
| 5 bp insertion | 5 bp insertion & 1 bp insertion | 50.0 | 105 | - | C | 48.5 |
| 10 bp insertion | 10 bp insertion & 1 bp insertion | 50.0 | 105 | - | C | 48.2 |
| 50 bp insertion | 50 bp insertion & 1 bp insertion | 50.0 |  |  | Failed |  |

SNPs (1-3 per read) and indels (5-50 bp length) at 50% frequency together with sequencing errors were simulated as the control reads group. Then a 1 bp insertion at position was inserted into the control group to form inserted read group. Co-analysis were conducted on both inserted reads and control reads. For all read types except the
50 bp insertion, CRISPR-detector successfully identified true editing events (the 1bp insertion) without false positives of variants existing in control samples.

CRISPResso2 also offers batch mode to output and visualize variants at the same location from different samples. While this allows the convenience of comparing more than 2 samples, the ability to directly filter germline and other existing variants from control samples is missing. In conclusion, CRISPR-detector significantly increases users’ efficiency in recognizing and confirming target variants.

#### 2. Scalability on Pooled Amplicons

In a few years, the scope of NGS-based on/off-target editing detection has expanded from only a few amplicons to hundreds of amplicons, to the coverage of all potential off-target sites, and even to WGS to ensure that every portion of the genome is screened. From a recent review, current mainstream in silico off-target prediction algorithms reach 95% sensitivity if the top 200 potential off-target sites are considered (Bao et al., 2021). This finding indicates a requirement for analysis tools to consider speed and scalability during development. To deal with this need, CRISPResso2 provides two specific pipelines, CRISPRessoPooled and CRISPRessoWGS, for these two input types. In comparison, CRISPR-detector’s optimized high-performance modules can handle small and large datasets from one single pipeline to provide uncompromised accuracy and scalability.

For benchmarking, we generated 200 amplicons in the format of 150 bp paired-end reads in which a large panel of SNPs, indels, and sequencing errors were introduced, similar to our single amplicon benchmark. Accuracy and speed performance are no different from the single amplicon test, and CRISPR-detector consistently outperforms CRISPResso2 (Fig. 2C and 2D). It should be noted that computation speed is not a bottleneck for small data. However, for larger datasets such as WGS, alignment and variant calling are two critical steps affecting pipeline speed, and CRISPR-detector utilizes and benefits from Sentieon’s high-performance alignment and sort modules.

#### 3. Whole Genome Sequencing Performance

The main difference between WGS and amplicon datasets are the regions of analysis (∼3 GB vs < 1 KB) and the sequencing depth (< 100x vs > 10,000x). Whole genome screening identifies more false positives, whereas insufficient sequencing depth makes false negatives unavoidable. To evaluate the accuracy of CRISPR-detector on a WGS analysis, we inserted simulated SNPs and indels into one chromosome sequence using BAMSurgeon (Ewing et al., 2015). Specifically, approximately 990 variants with 1-100% AFs were inserted into a 100x WGS dataset, and approximately 967 variants with 3.3-100% AFs were inserted into a 30x WGS dataset. To ensure at least 3 reads to support a reported variant, we only reported variants with AFs above 5% for 100x WGS and AFs above 15% for 30x WGS. Inserted reads and original reads are processed in co-analysis mode.

We found that co-analysis results were always better than only analyzing edited samples (Table 2A). The accuracy of SNPs is lower than that of amplicons, but acceptable, especially for the co-analysis of SNPs with AF > 15%. The accuracy of indels is significantly lower than that of amplicons, which is partially due to the fact that reads with indels are more difficult to map to the correct location in the full genome. Thus the reported indel frequency is lower than the correct number and may be filtered out by AF cutoff.

**Table 2.** SNP, indel, and SV Performance on WGS datasets.

| AF > 5% | SNP |  | Insertions (1-50 bp) |  | Deletions (1-72 bp) |  |
| --- | --- | --- | --- | --- | --- | --- |
|  | Sensitivity% | Specificity% | Sensitivity% | Specificity% | Sensitivity% | Specificity% |
| 100x co-analysis | 87.7 | 82.4 | 58.3 | 39.3 | 56.5 | 37 |
| 100x edited only | 96.4 | 41.4 | 55.6 | 13.9 | 71.7 | 18.3 |
| AF > 15% | SNP |  | Insertions (1-50 bp) |  | Deletions (1-72 bp) |  |
|  | Sensitivity% | Specificity% | Sensitivity% | Specificity% | Sensitivity% | Specificity% |
| 30x co-analysis | 98.8 | 99.3 | 13.3 | 50 | 23.7 | 66.7 |
| 30x edited only | 97.3 | 70.9 | 16.7 | 18.8 | 23.7 | 27.3 |

B
| Reads Type |  | 30x Short Reads | 100x Co-barcoded Reads |  | 100x Long Reads |
| --- | --- | --- | --- | --- | --- |
| SV Length | Pipeline | CRISPR-detector | Smoove | Long Ranger | Sniffles |
| 50-1000 bp | TP | 1268 | 724 | 2304 | 4168 |
|  | FP | 302 | 248 | 1279 | 5285 |
|  | FN | 3451 | 3995 | 2415 | 551 |
|  | Recall | 26.87% | 15.34% | 48.82% | 88.32% |
|  | Precision | 80.76% | 74.49% | 64.30% | 44.09% |
| > 1000 bp | TP | 445 | 463 | 558 | 570 |
|  | FP | 212 | 162 | 385 | 447 |
|  | FN | 172 | 154 | 59 | 47 |
|  | Recall | 72.12% | 75.04% | 90.44% | 92.38% |
|  | Precision | 67.73% | 74.08% | 59.17% | 56.05% |

**A: SNPs and indels accuracy benchmark.** SNPs and indels were introduced into one chromosome sequence. Only variants with AFs above 5% for 100x WGS, and with AFs above 15% for 30x WGS were reported to ensure at least 3 reads to support a reported variant. Sensitivity is calculated by $TP/(TP+FN)$ , and specificity is calculated by $TP/(TP+FP)$ [TP: True Positive; FN: False Negative; FP: False Negative]. Inserted reads were processed in both co-analysis mode and edited only mode. Co-analysis showed better accuracy than only analyzing the edited sample, especially for specificity%. Due to a much larger analysis region, the accuracy of WGS tends to be lower than that of amplicons. **B: SV accuracy benchmark.** We utilized the well-established NIST GIAB reference sample HG002 dataset to serve as a truth set. SVs were called from the CRISPR-detector pipeline and compared with the NIST truth set using Truvari to determine True Positive (TP), False Positive (FP), and
False Negative (FN), and then to calculate Recall and Precision values. We also displayed the results of other types of data and pipelines from a recently published study that used the same HG002 truth dataset (Zook et al., 2020). In the 50 bp - 1 kb group, CRISPR-detector reaches 26.87% recall and 80.76% precision and slightly outperforms Smoove. Long Ranger returned similar accuracy to CRISPR-detector, and Sniffles returns better recall than any other callers. For longer deletions (> 1 kb), CRISPR-detector's accuracy score is slightly lower than co-barcoded or long-read callers.

To validate this performance, a pair of 30x WGS samples (“Mutant” and “Rescued”) from a previous study was used (Wu et al., 2015). “Mutant” reads are from control cells having a pre-existing homozygous 1 bp disease causing deletion relative to the reference genome in the coding sequence of the mutant gene *Crygc*, “Rescued” means the edited clone rescued by CRISPR-Cas9-introduced 1-bp insertions 6-7 bp from the deletion. CRISPR-detector completed the “Mutant” group WGS dataset in 3 repeated runs using a consistent and reasonable 4h19m running time with 15 threads allocated (in total 65 core-hours), and it successfully detected the homozygous 1 bp deletion (chr1 65118239) as expected. Similarly, the pipeline completed the “Rescued” group sample in 5h36m and detected 2 insertions 6-7 bp from the deletion (chr1 65118232; chr1 65118233). CRISPRessoWGS was not included in this comparison, because in order to run WGS samples it requires a BED file, which is not defined by the original study.

### Structural Variation Identification

Gene editing introduces more than just short variants. For example, in viral-mediated gene transfer events, it is important to know the exact genome insertion location of the viral vectors to evaluate off-target event occurrence and functional consequences. Therefore, structural variations should be considered in the analysis pipeline design. A major purpose for conducting WGS on edited samples is to better identify off-target structural variants. To benchmark the accuracy, we designed two datasets for both general SV calling and vector insertion events.

We utilized the well-established PrecisionFDA dataset from HG002, in which a total of 12,745 isolated, sequence-resolved insertion (7,281) and deletion (5,464) calls longer than 50 bp are available, to serve as a truth set (Zook et al., 2020). SVs were called from the CRISPR-detector pipeline and compared with the NIST truth set using Truvari (https://github.com/spiralgenetics/truvari). To better evaluate the performance of CRISPR-detector and to better understand the limitations of short read-based SV calling compared to co-barcoded reads and long reads, we also compared the result with some pipelines from a recently published study that used the same testing dataset (Guo et al., 2021). Because insertions are more challenging and Truvari can only recognize the exact match to the truth set, which significantly underestimates true positives, we only showed the performance on deletions (Table 2B).

Deletions were grouped into two categories based on their length. In the 50 bp – 1 kb group where the majority of deletions exist, CRISPR-detector achieves 26.87% recall and 80.76% precision and slightly outperforms Smoove (https://brentp.github.io/post/smoove) on 100x co-barcoded reads. Long Ranger (https://github.com/10Xgenomics/longranger) returned similar accuracy to CRISPR-detector on 100x co-barcoded reads, and Sniffles returns better recall than any other callers (Sedlazeck et al., 2018). For longer deletions (>1k), CRISPR-detector’s accuracy score is slightly lower than co-barcoded or long-read callers. It is widely accepted that long read sequencing and co-barcoded read sequencing technologies are more capable of identifying large deletions, especially for improving recall, but the CRISPR-detector pipeline on short reads data still generates useful results. For the most accurate SV detection, long read whole genome sequencing is preferentially recommended, and CRISPR-detector will aim to adapt to these long read types in a future release.

Next, we simulated 150 bp paired-end reads with mimic ∼4.7k AAV vector insertion in a 30x WGS dataset to see if CRISPR-detector reports the insertion location. Because the inserted sequences are known, we modified the analysis pipeline to overcome the incompetence of insertion detection from the general SV calling pipeline. The AAV vector sequence was added into the human reference genome file as a dummy chromosome. Reads thus can be mapped to both human and vector sequences, and a translocation event can be reported to show the insertion location. We also introduced AAV partial sequences to better mimic partial insertion and simulated five scenarios, including insertion, reverse insertion, non-AAV sequence insertion, partial swap, and insertion with foreign sequences. In all 5 scenarios, CRISPR-detector returned the correct insertion location and always captured at least one of the breakpoints (Fig. S2), giving users the ability to take a closer look at the reported region and confirm the exact insertion format.

### Assessing Real World Clinical Data

In addition to simulated datasets, we utilized 3 published clinical datasets to evaluate the accuracy of CRISPR-detector and compared our analysis either with the original results published with the datasets or with CRISPResso2. The 3 datasets included are 1) rhAMP-seq from IDT; 2) ABE8e-NRCH (Newby et al., 2021); and 3) SPACE (Grünewald et al., 2020).

The first dataset benchmarked was the rhAMP-seq sample data provided on the IDT website. For the true editing positions provided by IDT (detected by GATK4 with a tuned parameter) (Poplin et al., 2018), CRISPR-detector achieved 100% accuracy with no false positive or false negative reports (Table S1).

CRISPR-detector is also suitable for assessing base editing outcomes. We analyzed 479 off-target sites from the third primer pool identified by Cas-OFFinder and CIRCLE-seq (Bae et al., 2014; Tsai et al., 2017) that were amplified from genomic DNA from CD34+ SCD cells edited by ABE8e-NRCH or from unedited control SCD donor cells using rhAMP-Seq (IDT). Following the original study’s methods, we calculated the statistical significance of ABE8e-NRCH mRNA off-target editing compared with control samples, by applying a Chi-square test. The 2×2 contingency table was constructed based on the number of edited reads and unedited reads, in treated and control groups. FDR was calculated using the Benjamini/Hochberg method. We applied the same cutoff thresholds as the original study: (1) FDR < 0.05 and (2) difference in editing frequency between treated and control > 0.5%.

As a result, among the 479 sites, 14 sites in the replicate 1 sample and 29 sites in the replicate 2 sample were identified as containing A•T-to-G•C mutations of significance. In replicate 1 sample, CRISPR-detector and CRISPResso2 agreed on all 479 sites. In the replicate 2 sample, the two pipelines showed discordance at only 3 sites (Fig. S3). Specifically, site “91469BDFFDAD44DZ0Z (off-target site 54)” was called positive only by CRISPResso2. The difference in editing frequency for this site between the treated and the control was 0.8%. Two sites were called positive only by CRISPR-detector. The differences in editing frequency between treated and control at these sites were near the 0.5% cut-off value. While the truth is not available to determine which calls are correct, current information indicates that these calls are on the edge between positive and negative (Table S2).

The dual-deaminase base editor SPACE (Synchronous Programmable Adenine and Cytosine Editor) can concurrently introduce A•T-to-G•C and C•G-to-T•A substitutions (Grünewald et al., 2020). CRISPR-detector’s performance was compared with CRISPResso2 for 28 SPACE target sites with nCas9 treatment as control. Each potential A•T-to-G•C and C•G-to-T•A position evaluated was categorized into 5 levels based on reported frequency: < 1%, 1%-10%, 10%-30%, 30%-60%, > 60%, and the highest frequency within the 20-bp evaluation window was chosen to represent the overall editing level for this protospacer. All 28 nCas9 samples showed no targeted substitution, but every SPACE treated sample returned multiple substitutions within the window.

CRISPR-detector and CRISPResso2 reported generally concordant editing levels for all 28 SPACE-treated sites (Fig. S4); only 10 of 560 (1.8%) evaluated positions showed discordant levels (Table S3). At these discordant positions, CRISPR-detector typically reported slightly lower frequencies than CRISPResso2, which was the same trend seen in our observations using simulated datasets.

In conclusion, CRISPR-detector showed identical accuracy as the IDT original analysis pipeline for the rhAMP-seq dataset and the same accuracy as CRISPResso2 on the ABE8e-NRCH and SPACE datasets.

### Functional Annotation of Editing Induced Mutations

ClinVar database is one of the largest public databases on human genomic variants and interpretations of their relationships to diseases and other conditions maintained by NCBI. Our pathogenicity annotations utilize this database but the outcome may differ from true pathogenicity. We examined the annotation results of the processed SPACE and ABE8e-NRCH datasets and found 1 variant returning annotation from the ClinVar database. From the SPACE datasets, a stopgain C•G-to- T•A mutation on *FANCF* site 1 was annotated to be pathogenic by ClinVar, and associated with Fanconi anemia. Our tool provides a convenient way to display ClinVar annotation if an off-target mutation exists in the ClinVar database.

We have added a clear statement in our user agreement that “The pathogenicity annotations are based on the ClinVar database.” The annotations are not intended for direct diagnostic use or medical decision-making purposes.

## Discussion

### Comparison with Other Analysis Tools

In pace with the development of genome engineering, analysis toolkits to assess the quality of genome editing have been developed and made available to the community. Early published tools such as CRISPR-GA (2014), ampliconDIVider (2015), and Cas-Analyzer (2017) only support single amplicon sequencing data, whereas AGEseq (2015) and CRISPResso2 (2019) support pooled amplicon sequencing data but do not support unbiased analysis of WGS data or SV calling. Several widely-used tools for analyzing genome editing NGS data were compared (Table 3). Clearly, CRISPR-detector is more comprehensive with functions that other listed toolkits are not capable of, including genetic background removal and mutation annotation, which significantly improve accuracy and show possible functional and clinical consequences.

**Table 3.**
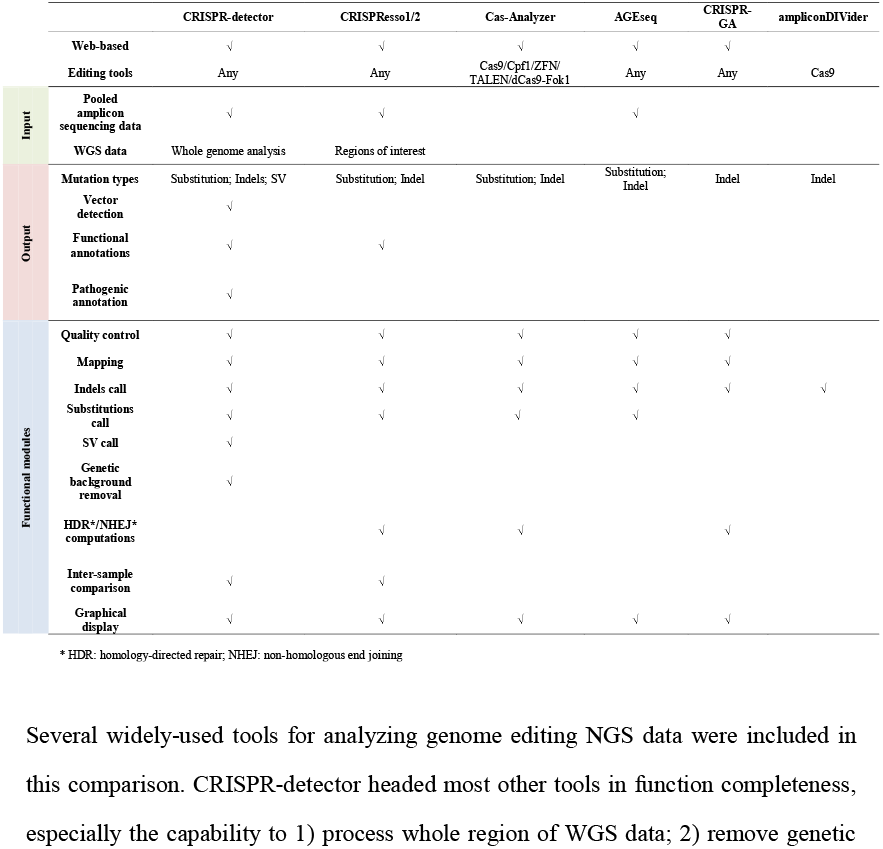
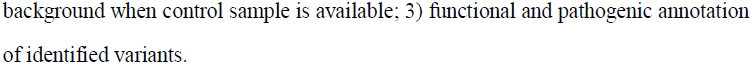
Comparing functions of tools that assess genome editing results.

### Other Potentially Useful Computation Modules

We considered whether read trimming and filtering steps should be included in the pipeline and decided they were unnecessary. Sequencing quality has improved greatly recently, and most modern sequencers are capable of generating raw reads with only rare bases below Q30, and thus by default, the read filtering step is not included. We suggest that users conduct QC using CRISPR-detector if they expect sequencing quality issues. We also decided to exclude adapter and primer trimming from the pipeline because primers are typically not included in the reference target regions in this application. Adapters may exist in raw data, but they do not affect the mapping result to a noticeable extent, and thus removing them may not be worth the computational cost.

### Benefits of Variant Clinical Annotation

Genomic variants may lead to clinically meaningful changes to protein function and phenotypes, and it is important for off-target detection to evaluate the potential clinical outcome of these introduced variants. The key here is to separate pathogenic variants from others. Instead of requiring users to take the variant output and conduct annotation on other platforms, we integrated the functional and clinical annotation within the CRISPR-detector platform so that users can directly assess potential clinical results.

## Materials and methods

### Pipeline Description

As shown in the CRISPR-detector flowchart (Fig. 1A), CRISPR-detector accepts single/pooled amplicon deep-sequencing as well as WGS data as input. Input reads are mapped to amplicon sequence(s) or a reference genome to call variants. If a control group sample is provided, CRISPR-detector will use it to remove background variants prior to analyzing the genome editing events of the treatment group sample.

For targeted amplicon sequencing data, basic information including amplicon sequence(s) is compulsory, whereas sgRNA sequence(s) input is optional. After variants are called, the CRISPR-detector web server supports the clinical annotation of mutations for human samples and functional annotation for nine other species. The locally deployable pipeline supports the use of a specific database built by users for mutation annotation. The CRISPR-detector generates graphs and tables that quantify and visualize the location and type of mutation(s) within the amplicon sequence(s).

For WGS data analysis, a reference genome is mandatory as input. The BED format text file is optional and is used for calling variants at regions of interest such as on/off-target sites. Our CRISPR-detector tool is capable of calling variants involving SNPs and SVs. CRISPR-detector outputs a VCF (Variant Call Format) file, which contains information about the variants called. Efficient tools for in vivo gene transfer, such as AAV viral vectors (rAAVs), may nonspecifically integrate into the host cell genome and cause insertional mutagenesis events, and if the viral vector sequence(s) is provided, CRISPR-detector can also report the location of vector insertion events.

### Webpage Input Module

On the CISPR-detector home page (https://db.cngb.org/crispr-detector), users can upload FASTQ files, provide basic information, and set parameters. FASTQ sequences from the treatment group are mandatory, whereas sequences from the control group are optional, but recommended. Each of the input FASTQ files can be sequencing data from a single amplicon or pooled amplicons. Although CRISPR-detector does not provide sample separation based on sample indexes or barcodes, it does support separated FASTQ files from pooled amplicon sequencing protocols similar to those described in Hi-TOM (Liu et al., 2019).

### Data Analysis Module

CRISPR-detector analyzes the input data in two stages: “preprocessing” and “variant identification”. In the preprocessing stage, reads from edited samples or control samples are first mapped to either amplicon sequences or the human reference genome using Sentieon Minimap2 (Li, 2018), which is an accelerated version of the corresponding open-source Minimap2 that returns identical results. Next, all mapped reads are sorted by the Sentieon “sort” utility when read ranks are adjusted by their genomic locations. Then, a quality control step is conducted on the sorted BAM files by the Sentieon “Quality Yield” utility to calculate a quality score distribution of the aligned reads. A warning will be given to the users if the Q30 value (percentage of mapped bases with quality scores greater than 30) is lower than 0.75 by default. The claimed Q30 for Miseq is ∼0.85, and thus a Q30 value lower than 0.75 may indicate abnormal sequencing quality issues. The last step of this stage is deduplication by the Sentieon “Dedup and LocusCollector” utility, which marks reads with the same starting and ending positions to avoid duplicate counting. This “dedup” step is necessary for WGS or probe hybridization-based capture sequencing but should be skipped for amplicon sequencing.

The second “Variant Identification” stage transforms prepared BAM files into variants. To begin, Sentieon TNscope is used with the default parameters to generate SNP and indel candidate variants. Sentieon TNscope is developed on top of GATK Mutect2 v3.8 and is specifically optimized for low AFs variant calling and integrated structural variation calling functions (Freed et al., 2018; Benjamin et al., 2019). Then structural variations are called by TNscope using default parameters if the pipeline is run as a local version. Genetic background filtering is the next step, in which TNscope generates candidate variants from control samples. Each candidate variant from edited and control samples is tested with Fisher’s exact test to calculate a p-value and determine if its supporting features reach significance. Variants from edited samples are filtered out if the p-value is higher than the user-determined cutoff (default 0.05). The next step is to filter out low-quality reads. Specifically, reads marked by TNscope other than “PASS”, “triallelic_site”, or “alt_allele_in_normal” are filtered out, and variants with lower than input AF cutoff (default 0.005) or with less than input supporting read depth (default 500) for amplicon data are filtered out. The last step is to filter out variants that are not likely to be introduced by editing. When uploading an amplicon dataset, users can provide guide RNA sequence(s) and input a “cleavage_offset” parameter to define the center of the quantification window with respect to the 3’ end of the provided sgRNA sequence excluding the PAM. The “window_size” parameter can also be designated to define the size (in bp) of the quantification window extending from the center of the quantification window defined by the “cleavage_offset” parameter. The default value of “window_size” is 0, meaning analysis is conducted for whole amplicons. If the quantification window is defined then any variants that do not overlap with the window are filtered out.

Functional and clinically important variants in the treatment group data are annotated using the published ANNOVAR (version 2019 Oct24) tool and ClinVar (version 20210501) databases (Wang et al., 2010). Functional variants provided by ANNOVAR include frameshift, nonsynonymous, start-gain/loss, or stop-gain/loss mutations (Fig.S5), and clinical variants are those marked pathogenic or likely pathogenic in the ClinVar database, which provides the interpretation of variant correlation with disease.

### Result Visualization Module

By clicking the “Example” button on the webpage, example results for real datasets are shown. The results webpage includes three parts:

1. A downloadable table summarizing the number of processed reads and the frequency of different types of mutations on all the input loci (Table S4).
2. A batch output mode comparing the editing outcomes from all input loci, producing figures comparing the general SNPs and indel ratios.
3. Results for individual input locus. The example figure shows the edited number position of the identified SNPs and indels (Fig. 3A) and the indel size distribution (Fig. 3B). The reported positions are from the co-analysis if control sample data are available. Furthermore, detailed information on mutations including the annotation of this locus is summarized in a downloadable table (Fig. 3C).

**Fig. 3.**
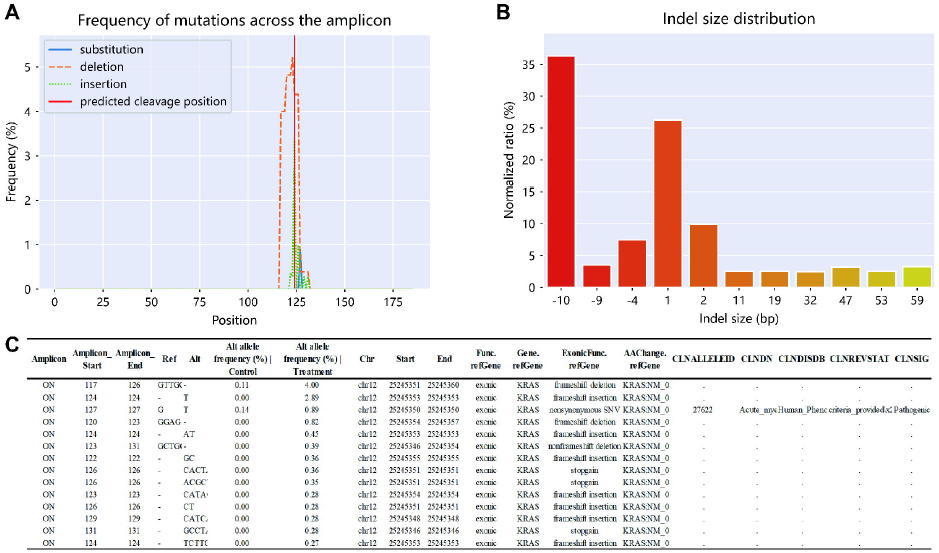
Example of data visualization for a single locus. These three figures and tables are part of the visualized result of the web-hosted version of CRISPR-detector pipeline. **A**: Frequency of identified SNPs and indels across the amplicon. **B:** Indel size distribution. **C:** Detailed information on identified variants on this locus, including function and clinical annotations.

### Locally Deployable Version

Due to data upload speed limitations, our web-hosted pipeline is inconvenient for processing WGS data. To provide an alternative solution, a version for local deployment that accepts NGS data larger than 100 MB, including WGS sequencing data, has also been developed and is provided to the community (available at https://github.com/hlcas/CRISPR-detector).

There are three differences between the web version and this locally deployable version: 1) the local pipelines accept WGS and other large datasets; 2) structural variants can be called; and 3) visualization modules and annotation functions are not included, because the pipeline outputs VCF format files.

Structural variations are called using TNscope, which is based on evidence from soft-clipped mapped reads and the discordance of read pairs. Specifically, the calling occurs in three stages. In the first stage, the caller identifies all abnormally aligned reads from treatment and control samples (if available). Concurrently, it builds a local assembly around the split reads and filters out all haplotypes that exist in the control samples and noisy haplotypes with little read support. In the second stage, it analyzes the candidate haplotypes assembled around split reads and identifies consistent haplotype pairs. These reads represent potential structural variant break-ends. In the third stage, the caller looks for evidence from the discordant read pairs.

## Data availability

Source code and scripts to simulate sequencing data are available at GitHub (https://github.com/hlcas/CRISPR-detector). WGS datasets are available as the GEO reference series SRP045395. SPACE and ABE8e-NRCH datasets can be accessed at the NCBI Sequence Read Archive (PRJNA609075; PRJNA725249).

## Credit authorship contribution statement

**Lei Huang, Haodong Chen, Jinnan Hu**: Conceptualization, Data curation, Formal analysis, Investigation, Methodology, Resources, Software, Validation, Visualization, Writing – Original draft, Review & Editing. **Xuechen Dai, Chuan Liu, Anduo Li, Xuechun Shen, Chen Qi**: Investigation, Methodology, Validation, Writing – Review & Editing. **Haixi Sun, Dengwei Zhang**: Conceptualization, Methodology, Supervision, Writing – Review & Editing. **Yuan Jiang, Dan Wang, Tong Chen**: Conceptualization, Funding acquisition, Investigation, Methodology, Project administration, Supervision, Writing – Original draft, Review & Editing. All authors read and approved the final manuscript.

## Acknowledgments

We would like to thank Dr. Gao Qianqian for advising on our web server and providing sequencing data and colleagues at China National GeneBank (Shenzhen) and EHBIO Gene Technology (Beijing) for their guidance and help in constructing the web server. The authors declare that the research was conducted in the absence of any commercial or financial relationships that could be construed as a potential conflict of interest. This work was supported by the Fundamental Research Funds for the Central Public Welfare Research Institutes (ZZ13-YQ-095 and ZZXT201708) and the Start-up Research Fund from BNU-HKBU United International College (UICR0700053-23).

