## Supplementary Data for "CRISPR-Detector: Fast and Accurate Detection, Visualization, and Annotation of Genome-Wide Mutations Induced by Gene Editing Events"

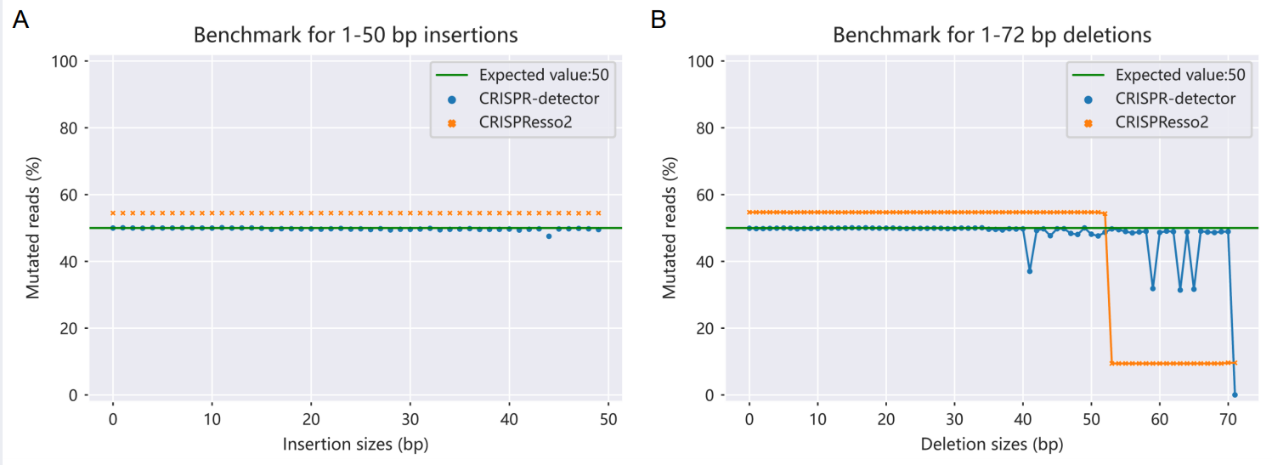

**Fig. S1. Benchmark for 1-50 bp insertions and 1-72 bp deletions. A:** Both CRISPR-detector and CRISPResso2 correctly report 1-50 bp insertions, while CRISPResso2’s reported value was higher than expected. **B:** CRISPR-detector failed to report insertions larger than 72 bp while CRISPResso2 failed to report insertions larger than 53 bp. The waves are likely caused by the randomness of the deletion location relative to read ends. Reads will not be able to be correctly mapped if the deletion position is close to the ends of the reads.

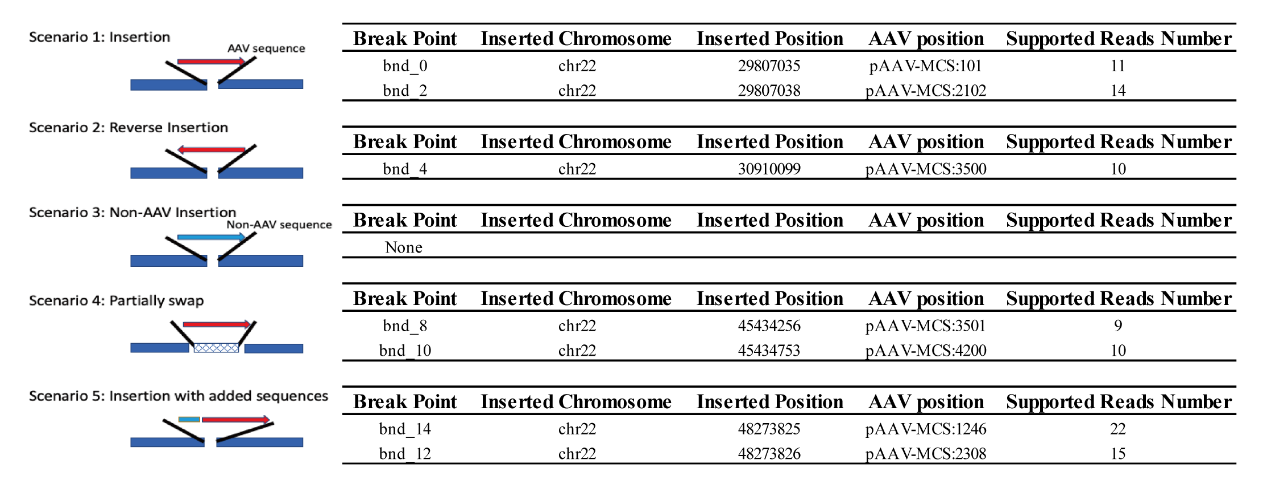

**Fig. S2.** **AAV sequence insertion detection.** 150 bp paired-end reads were generated to mimic ~4.7k AAV vector insertion in a 30x WGS dataset. Before reads alignment, the AAV vector sequence was added into the human reference genome file as a dummy chromosome. Reads thus can be aligned to both human and vector sequences, and a translocation event can be reported if part of a read pair is mapped to vector sequences and the other part is mapped to the human genome. Five simulated scenarios were tested, including insertion, reverse insertion, non-AAV sequence insertion, partial swap, and insertion with foreign sequences. CRISPR-detector returned the correct insertion location in all five scenarios.

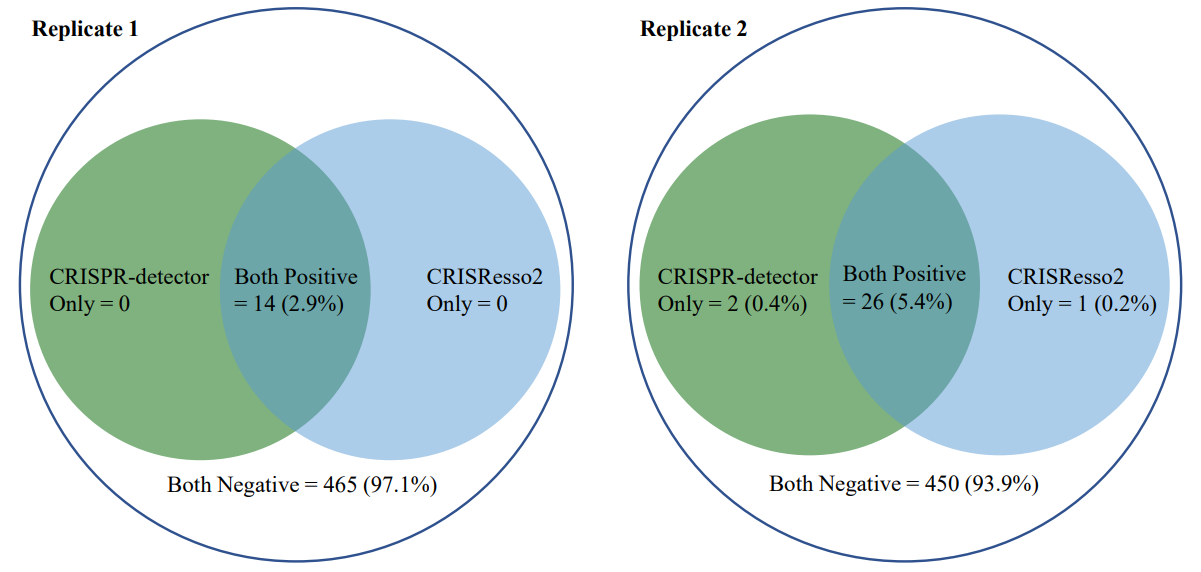

**Fig. S3.** **Venn diagram of CRISPR-detector and CRISPResso2 calling on ABE8e-NRCH datasets.** In replicate 1 sample, CRISPR-detector and CRISPResso2 agreed on all 479 sites for off-target events. And in the replicate 2 sample, the two pipelines showed discordances on only 3 sites. Most sites were either “Both Positive” (called detectable by both pipelines), or “Both Negative”. Positive sites were called based on: (1) FDR < 0.05 and (2) the difference in editing frequency between treated and control > 0.5% (Table S5).

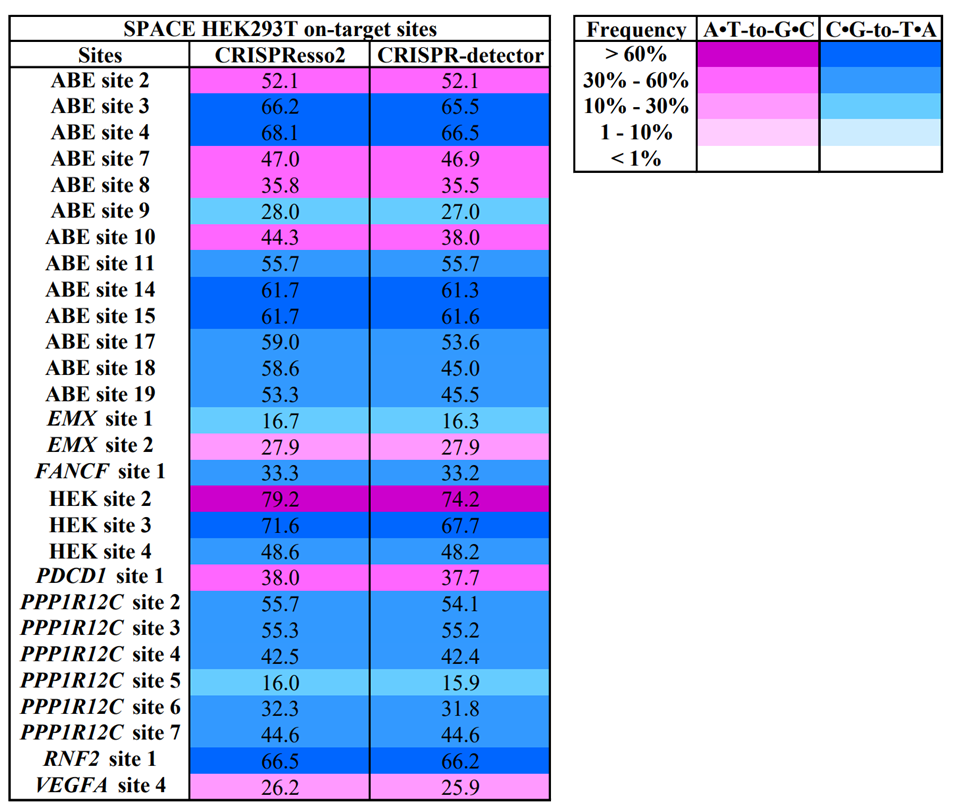

**Fig. S4. Maximum editing frequency across the protospacer.** CRISPR-detector’s performance was compared with CRISPResso2 on 28 on-target sites processed by SPACE. Each potential A•T-to-G•C and C•G-to-T•A position evaluated was color coded into 5 levels based on reported frequency: < 1%, 1%-10%, 10%-30%, 30%-60%, > 60%, and the highest frequency within the 20 bp evaluation window was chosen to represent overall editing level for this protospacer. CRISPR-detector and CRISPResso2 reported concordance editing levels on all 28 SPACE treated sites.

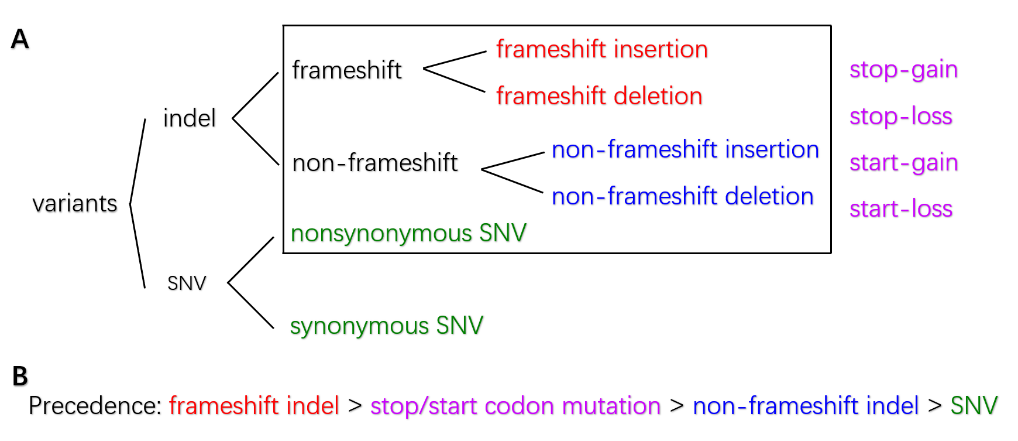

**Fig. S5. Functional consequences of exonic variants. A:** Functional consequences of the exonic variants could be annotated by ANNOVAR (Wang et al., 2010) as nonsynonymous SNV, synonymous SNV, frameshift insertion, frameshift deletion and non-frameshift insertion, non-frameshift deletion. Except for synonymous SNV, other types of exonic variants in the black border rectangle may cause stop-gain, stop-loss, start-gain, or start-loss. **B:** The precedence that ANNOVAR takes to decide what consequences to print out when a variant fits multiple categories.
